# Venous Thromboembolism Risk in Head and Neck Cancer: Significance of the Preoperative Platelet-to-Lymphocyte Ratio

**DOI:** 10.1101/217042

**Authors:** Tristan Tham, Lauren Rahman, Christina Persaud, Peter Costantino

## Abstract

**Objective:** We aimed to investigate the association between the preoperative platelet-to-lymphocyte ratio (PLR) and venous thromboembolism (VTE) in head and neck cancer (HNC) patients undergoing major surgery.

**Study Design:** Retrospective cohort study

**Setting:** Academic tertiary hospital from 2011 to 2017

**Subjects and Methods:** Patients with confirmed HNC undergoing major surgery were included in this study. The preoperative PLR was recorded for all patients. Known VTE risk factors, including age, sex, smoking, BMI, prior VTE, and anticoagulation were also recorded. Risk factors were screened in univariate analysis using Wilcoxon’s rank sum test and χ2 test (Bonferroni corrected). Significant covariates were subsequently included in a multivariate regression model. Bootstrap techniques were used to obtain credible confidence intervals (CI).

**Results:** There were 306 patients enrolled with 7 cases of VTE (6 DVTs and 1 PE). On univariate analysis, length of stay (p = 0.0026), length of surgery (p = 0.0029), and PLR (p = 0.0002) were founded to have significant associations with VTE. A Receiver Operator Characteristic (ROC) curve was constructed, that yielded an AUROC of 0.905 (95% CI: 0.82 - 0.98). Using an optimized cutoff, the multivariate model showed that length of surgery (β 95% CI: 0.0001 - 0.0006; p = 0.0056), and PLR (β 95% CI: 5.3256 - 5.3868; p < 0.0001) were significant independent predictors of VTE.

**Conclusion:** This exploratory pilot study has shown that PLR offers a potentially accurate risk stratification measure as an adjunct to current tools in VTE risk prediction, without additional cost to health systems.

**Oral Presentation:** This data was presented as an oral presentation at the Annual American Academy of Otolaryngology – Head and Neck Surgery (AAO-HNSF) Meeting, 13th September 2017

## Introduction

Venous thromboembolism (VTE) is an uncommon but potentially life-threatening complication in patients with head and neck cancer (HNC) undergoing surgical resection. VTE as a disease entity includes both deep vein thrombosis (DVT) and pulmonary embolism (PE). Although the incidence of VTE in the general otolaryngology population is low, ^1–5^ the incidence in patients with cancer has been reported to be higher at 1.4% - 5.8%. ^5, 6^ This higher incidence rate of VTE in HNC patients is likely due to older age, tobacco use, major surgery, reduced mobility, and malignancy itself.^7^

Current methods of predicting VTE risk in HNC patients, such as the widely used Caprini VTE risk assessment ^8^, lose their discriminatory power within groups of high-risk patients. According to the Caprini assessment system, patients with HNC undergoing major surgery are at the minimum in the high-risk category. Previously, Thai and colleagues had found that at or above the highest Caprini risk level, the Caprini scoring system failed to discriminate between patients who were likely to develop VTEs.^6^ Similar conclusions were also obtained in a prospective study of VTE in HNC patients by Clayburgh et al.^9^ Therefore, there is a need to develop better VTE risk assessment tools that would assist head & neck surgeons in better stratifying their high-risk patients.

Within the cardiovascular and thrombosis research literature, there has been growing interest in utilizing the platelet-to-lymphocyte ratio (PLR) to predict thrombosis and mortality.^10–13^ The exact mechanism of how the PLR biomarker reflects intrinsic thrombotic propensity has not been fully elucidated, but it has been hypothesized that it is likely secondary to the complex interplay between intrinsic thrombotic (platelet count) and systemic inflammatory (lymphocyte count) factors. ^10–13^

In two recent studies, the PLR biomarker has been demonstrated to predict VTE risk in cancer patients.^12, 13^ These studies were performed in heterogeneous populations with different tumor types and treatments. They were also performed in the ambulatory setting, which could arguably be lower-risk setting than in the inpatient postoperative setting because of the patient’s mobility status. To the best of our knowledge, there have been no studies investigating the effect of the PLR in predicting VTE in HNC or in the surgical setting.

In this context, we performed this exploratory pilot study with the aim of investigating the predictive association of the pre-operative PLR for VTE in a high-risk cohort: HNC patients undergoing surgical resection.

## Methods

### Subjects

This retrospective cohort study included patients with HNC treated at the New York Head & Neck Institute (NYHNI) - Northwell Health System between 1^st^ January 2011 and 30^th^ June 2017. This study was approved by the Institutional Review Board (IRB) of the Northwell Health System under IRB# 17-0068-LHH. Because of the inability of current methods to stratify within high-risk HNC patients,^6, 9^ this study was designed to specifically only include patients who were at minimum ‘high-risk’ as determined by the Caprini scoring system,^8^ and others.^14, 15^ The inclusion criteria was (a) any patient with histologically confirmed malignant disease of the head and neck that is primary or recurrent, (b) undergoing major surgical resection. These patients are automatically classified as ‘high risk’ for VTE regardless of other factors such as comorbidities, length of immobilization, and age.^8^ The exclusion criteria for our study was (a) lymphoproliferative malignancies, (b) metastatic disease or palliative procedures, (c) minor procedures lasting < 1 hour, including but not limited to biopsies and tracheostomies, (d) incomplete medical records, including absence of preoperative complete blood count (CBC) differential, (e) CBC that was taken more than two weeks pre-operatively. Patients were screened using pre-defined ICD-9/10 diagnosis codes (Supplementary materials). All patients were treated according to the NYHNI post-operative protocol that included sequential compression devices (SCD) and early mobilization. Post-operative VTE prophylaxis was not mandated for all patients, and was administered according to the surgeon’s clinical judgement.

### Variable selection

The literature was reviewed for important VTE risk factors in HNC that could be used to build univariate and multivariate models.^5, 6, 9^ All established risk factors for VTE were included in the data collection process: Age, Sex, Smoking Status, body mass index (BMI), Length of Stay, Length of Surgery, Prior VTE, Postoperative heparin, Postoperative aspirin. Because of the expected low incidence of VTE, we decided *a priori* to exclude other variables (ambulation time, comorbidities) that had been shown to be insignificant in this population by previous authors, ^6, 9^ so as to avoid the multiplicity effect in the statistical model. ^16^ Additionally, we did not include scoring systems such as the Caprini system, Karnofsky performance status or Charlson comorbidity index, as none of them have been shown to be able to stratify VTE in high-risk HNC patients. ^6, 9^

### Data collection

The electronic healthcare record of patients was systematically reviewed, and data was stored in REDCAP (Research Electronic Data Capture), a secure, web-based data capture application hosted at the Northwell Healthcare System. ^17^ Demographic and clinical information was retrieved from the patient’s history and physical notes, operative reports, laboratory reports, progress notes, imaging reports, discharge summaries, medication order forms, outpatient notes, and anesthesia notes. Imaging reports for all patients were examined to find out which patients had received ultrasound Doppler or spiral CT. These reports were then accessed to confirm the presence of VTE. Preoperative lab results were retrieved to obtain the PLR and other CBC parameters. The PLR was calculated as the platelet count (10^3 / uL) divided by the lymphocyte count (10^3 / uL), both taken from the same blood sample, within two weeks of surgery. For additional descriptive data, pathology reports were also examined to determine the tumor site and histopathology.

### Endpoint

The primary endpoint of this study was VTE defined as any deep venous thrombosis (DVT) or pulmonary embolism (PE). VTE was confirmed by diagnostic imaging, with either doppler ultrasonography for DVT, or with spiral CT for PE. Only VTEs in the immediate postoperative period (during inpatient stay) were included.

### Statistical analysis

Because of the expected sparse data set, we didn’t impose the assumption of a Gaussian distribution and therefore used non-parametric statistical tests wherever possible. Continuous variables were expressed as the means ± standard deviations, and cases (VTE) were compared to controls (non-VTE) using Wilcoxon’s rank sum test. On the other hand, categorical variables are presented as number of patients in each group, and compared using the χ^2^ test. In order to compensate for multiplicity in the univariate analysis, we applied the Bonferroni correction to the α level. ^18^ After correction, a p value of < 0.0038 was considered significant in the univariate analysis. For all other analyses, a p value of < 0.05 was considered significant. To further investigate the PLR, we also constructed a boxplot to visualize the differences in both groups. The cutoff between a ‘high’ or ‘low’ PLR has not yet been unified in the literature. ^12, 13^ Therefore, we used a simple linear discriminant analysis (LDA) classifier between PLR and VTE to generate a receiver operator characteristic (ROC) curve. This ROC curve was used to obtain an optimal cutoff point for the PLR, and this cutoff point was chosen to maximize both sensitivity and specificity. A dichotomized PLR variable based on this cutoff was used to separate patients, and compared with a χ^2^ test. The multiple logistic regression model was performed using the ROC cutoff and other significant variables discovered in univariate analysis. In the inference stage, we used Firth’s penalized method for deriving the maximum likelihood estimators (MLE) to avoid serious bias due to the sparse events data.^19^ Firth’s method uses the Jeffrey’s prior and is used in many related applications. ^19–21^

We also utilized the “bootstrap method” to obtain a credible confidence intervals (CI), ^22^ which is indicated for uncertain parameter distribution in small samples.^23^ Very briefly, bootstrap resampling works on the premise that the data collected is the best available approximation of the true probability distribution of the actual data distribution (distribution for ∞ patients). Bootstrap resampling works by refitting many pseudo data sets. The pseudo data set is composed of the dependent and independent variables that are randomly selected from the original data set. Fitting is then carried out on these pseudo data sets. From this, bootstrap is able to quantitatively estimate how the original data might have varied. For each model uncertainty calculation, either 1,000 or 10,000 repeat fits were used, which was computationally demanding. Regression coefficients (β) and their 95% confidence intervals (CI) are presented. All statistical analyses were conducted using the R software, version 3.3.3 (R Development Core Team, Vienna, Austria). ^24^

## Results

### Descriptive characteristics

There were a total of 306 patients with HNC undergoing major surgery that were included this study between January 2011 and June 2017. Among these patients, there were 7 cases of VTE (6 DVTs, 1 PE) and 299 patients without VTE. This incidence rate of ~ 2.3% is in agreement with previous reports.^6, 25, 26^ The average age was 61.8 ± 14.4 years, and the majority of them were male (66.8%). The most common histological type was Squamous cell carcinoma (66.7%). The subsites were the Lip and Oral Cavity (72, 23.5%), Tonsil/Oropharynx (43, 14.0%), Larynx (44, 14.3%), Salivary Gland (42, 13.7%), Nasal Cavity and Paranasal Sinuses (28, 9.1%), Cervical Lymph Node and Unknown Primary (30, 9.8%), Skull base (22, 7.2%), Nasopharynx (6, 2.0%), Hypopharynx (2, 0.7%), Cutaneous Head and Neck (14, 4.6%), Other Sites (4, 1.3%). The mean platelet count, WBC count, lymphocyte count, and PLR were 232.46 ± 71.71, 7.02 ± 2.23, 1.68 ± 0.73, and 160.90 ± 83.33 respectively. Four patients had a prior history of VTE, but none of them developed postoperative VTE, thus the ‘Prior VTE’ variable was discarded from further analysis. Baseline clinical characteristics and laboratory parameters between the two groups are presented in Table 1.

**Table 1.**
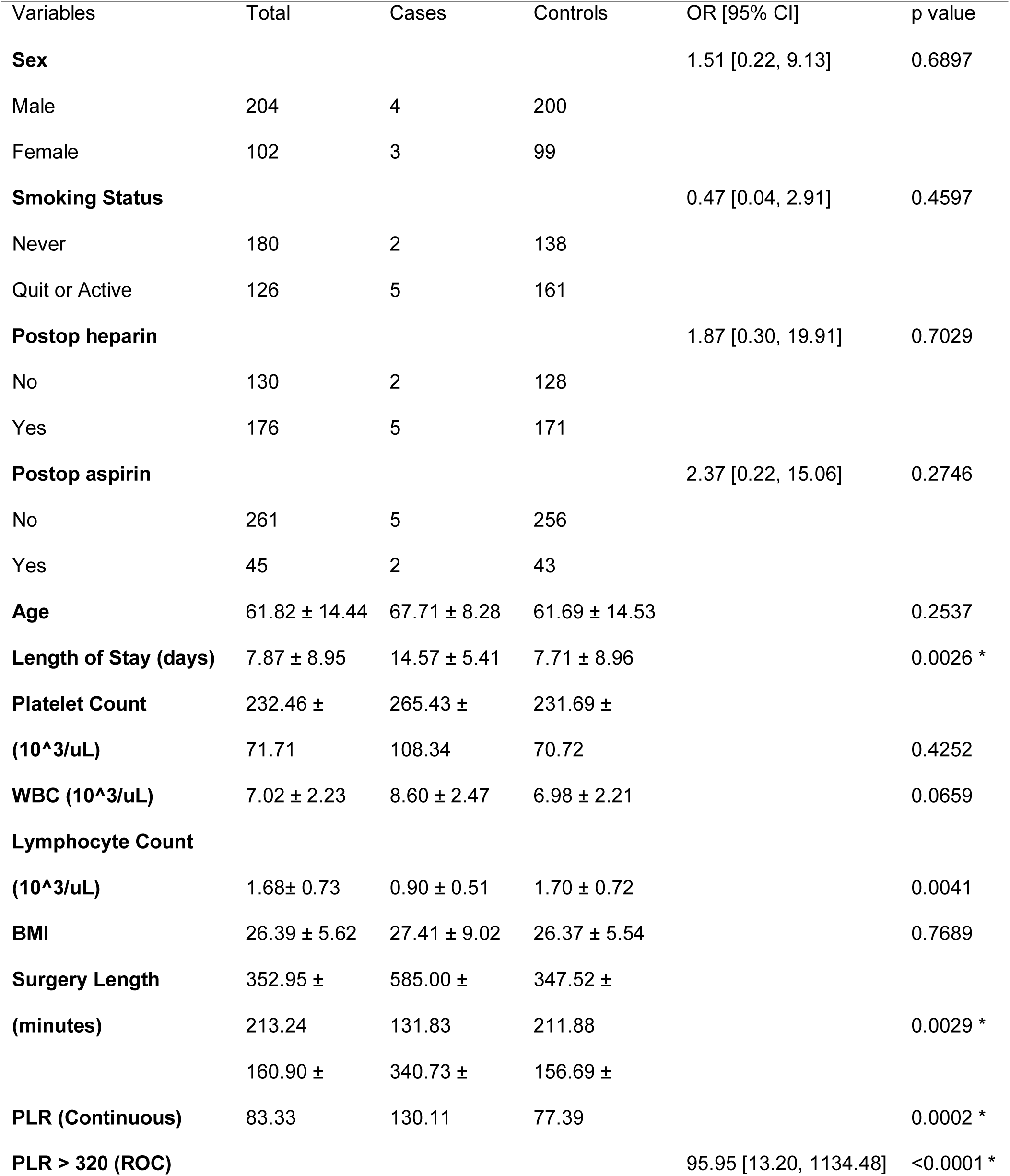
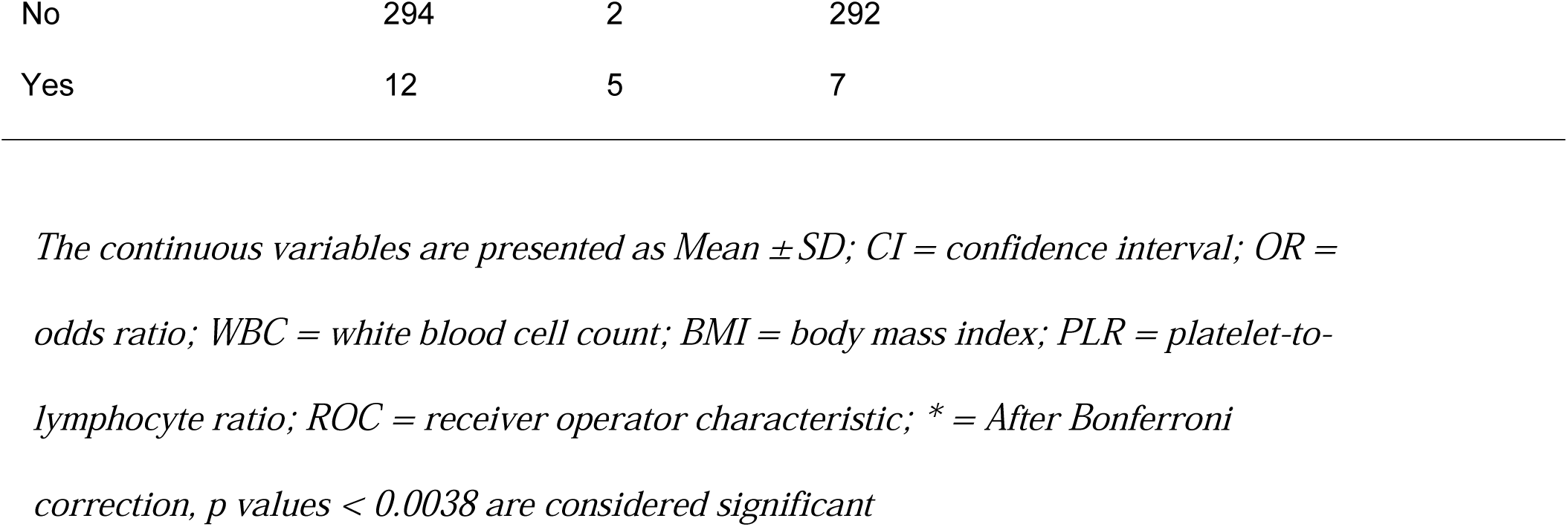
Patient Characteristics and Exploratory Univariate Analysis

| Variables | Total | Cases | Controls | OR [95% CI] | p value |
| --- | --- | --- | --- | --- | --- |
| **Sex** |  |  |  | 1.51 [0.22, 9.13] | 0.6897 |
| Male | 204 | 4 | 200 |  |  |
| Female | 102 | 3 | 99 |  |  |
| **Smoking Status** |  |  |  | 0.47 [0.04, 2.91] | 0.4597 |
| Never | 180 | 2 | 138 |  |  |
| Quit or Active | 126 | 5 | 161 |  |  |
| **Postop heparin** |  |  |  | 1.87 [0.30, 19.91] | 0.7029 |
| No | 130 | 2 | 128 |  |  |
| Yes | 176 | 5 | 171 |  |  |
| **Postop aspirin** |  |  |  | 2.37 [0.22, 15.06] | 0.2746 |
| No | 261 | 5 | 256 |  |  |
| Yes | 45 | 2 | 43 |  |  |
| **Age** | 61.82 ± 14.44 | 67.71 ± 8.28 | 61.69 ± 14.53 |  | 0.2537 |
| **Length of Stay (days)** | 7.87 ± 8.95 | 14.57 ± 5.41 | 7.71 ± 8.96 |  | 0.0026 * |
| **Platelet Count** | 232.46 ± | 265.43 ± | 231.69 ± |  |  |
| **(10<sup>3</sup>/uL)** | 71.71 | 108.34 | 70.72 |  | 0.4252 |
| **WBC (10<sup>3</sup>/uL)** | 7.02 ± 2.23 | 8.60 ± 2.47 | 6.98 ± 2.21 |  | 0.0659 |
| **Lymphocyte Count** |  |  |  |  |  |
| **(10<sup>3</sup>/uL)** | 1.68± 0.73 | 0.90 ± 0.51 | 1.70 ± 0.72 |  | 0.0041 |
| **BMI** | 26.39 ± 5.62 | 27.41 ± 9.02 | 26.37 ± 5.54 |  | 0.7689 |
| **Surgery Length** | 352.95 ± | 585.00 ± | 347.52 ± |  |  |
| **(minutes)** | 213.24 | 131.83 | 211.88 |  | 0.0029 * |
|  | 160.90 ± | 340.73 ± | 156.69 ± |  |  |
| **PLR (Continuous)** | 83.33 | 130.11 | 77.39 |  | 0.0002 * |
| **PLR > 320 (ROC)** |  |  |  | 95.95 [13.20, 1134.48] | <0.0001 * |

|  |  |  |  |
| --- | --- | --- | --- |
| No | 294 | 2 | 292 |
| Yes | 12 | 5 | 7 |

*The continuous variables are presented as Mean ± SD; CI = confidence interval; OR =* *odds ratio; WBC = white blood cell count; BMI = body mass index; PLR = platelet-to-* *lymphocyte ratio; ROC = receiver operator characteristic; \* = After Bonferroni* *correction, p values < 0.0038 are considered significant*

### Univariate and Multivariate Analysis

After application of the Bonferroni correction, variables Length of Stay (p = 0.0026), Surgery Length (p = 0.0029), and PLR (p = 0.0002) were founded to be significant in the univariate analysis. (Table 1) To further investigate the PLR, we constructed a boxplot to visualize the difference between cases and controls. (Figure 1) On visual inspection of the box plot, there was a difference between these two distributions, as suggested by previous literature. ^12, 13^ To obtain the PLR cutoff, an ROC curve was generated for VTE detection using PLR. (Figure 2) Using the bootstrap method, the AUC of the ROC model was ~ 0.91 (95% CI: 0.82 - 0.98), which is an indication of excellent discriminatory ability. ^27^ We then used the ROC curve to obtain the optimum cutoff point of 320, which had a sensitivity of 97.66% and specificity of 71.43%. Once the PLR was dichotomized to this cutoff level, a χ^2^ test was performed which showed significance (Table 1, p < 0.0001). Multiple logistic regression with 1,000 bootstrap iterations was performed for the significant variables discovered in the univariate analysis: PLR (ROC Cutoff), Length of Surgery, and Length of Stay. In the multivariate model, surgery length (p = 0.0056), and PLR (p < 0.0001) were found to be significant independent predictors of VTE. (Table 2) Of note, the regression coefficient (β) for PLR (95% CI: 5.3256 - 5.3868) was shown to be much larger than length of surgery (95% CI: 0.0001 - 0.0006).

**Table 2.** Multiple Logistic Regression with VTE, controlling for selected risk factors

| Variables | 95% CI with 1,000 Bootstrap iterations* | p value |
| --- | --- | --- |
| Surgery Length | [0.0001, 0.0006] | 0.0057^ |
| Length of Stay | [-0.0035, 0.0073] | 0.2922 |
| PLR > 320 | [5.3256, 5.3868] | <0.0001^ |

\* = Represents the 95% confidence interval of the regression coefficient ( $\beta$ ); ^ = p-values <0.05 are considered significant; CI = confidence interval; PLR = platelet-to-lymphocyte ratio

**Figure 1.**
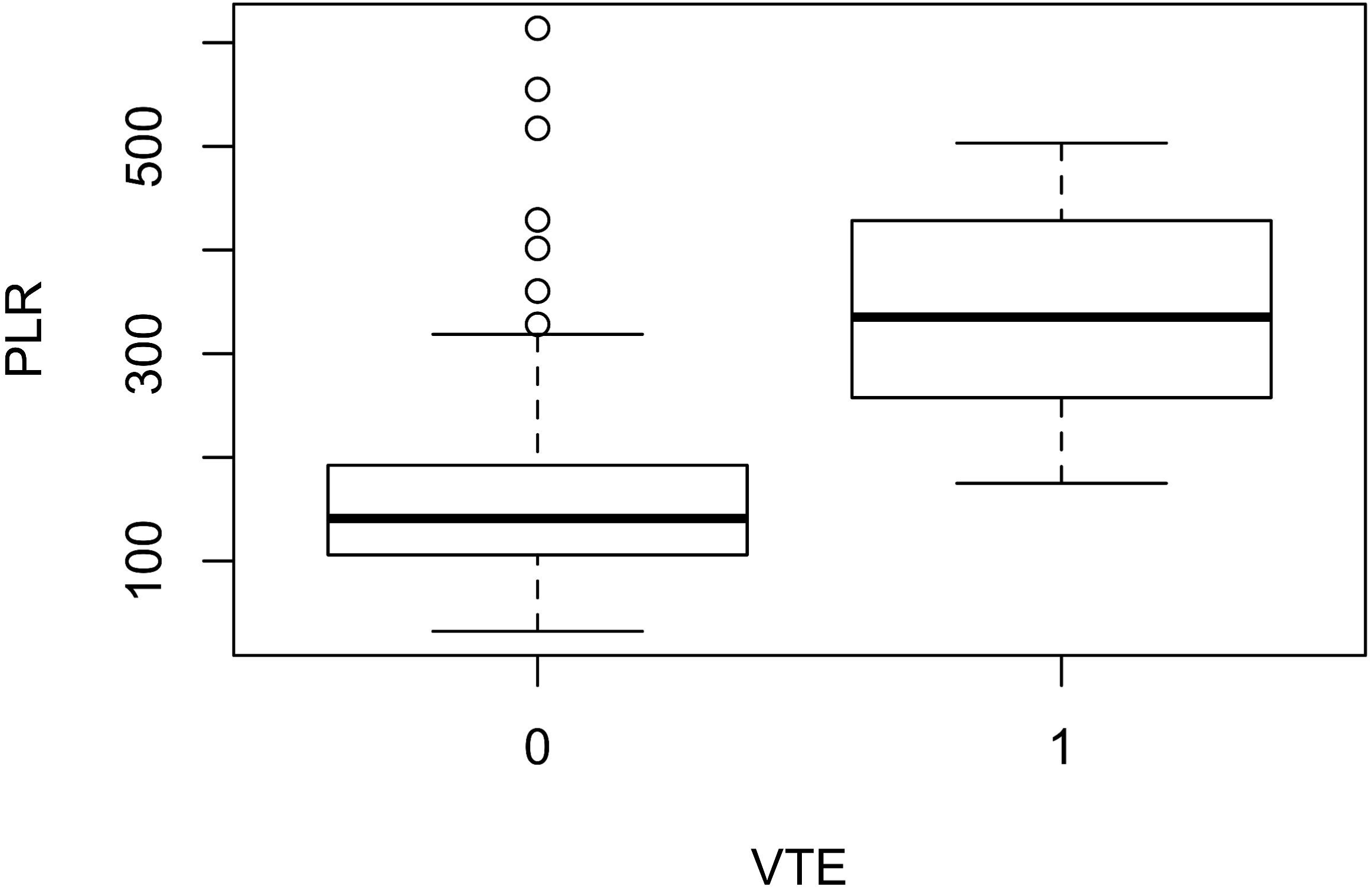
Boxplot of PLR vs VTE. This is a boxplot of the PLR values (y-axis) and presence of VTE (x-axis). Thick horizontal lines are medians, boxes show the interquartile range (IQR), error bars show the full range excluding outliers (circles) defined as being more than ± 1.5IQR outside the box.

**Figure 2.**
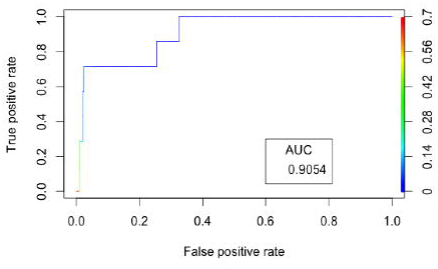
ROC Curve of VTE Detection Using PLR. ROC Curve for detecting VTE using the PLR value. This ROC curve was produced through 10,000 iterations of the bootstrap technique. The true positive rate (sensitivity) is plotted as a function of the false positive rate (1-specificity) for different positivity threshold (ie, different cutoff levels). The area under the ROC (AUROC) is 0.905 (95% CI: 0.82 - 0.98). The AUC is a measure of how well a quantitative test can distinguish between subjects with and without a disease.

## Discussion

In this study, we retrospectively analyzed the clinical and laboratory data in HNC patients undergoing major surgery, comparing patients with and without VTE. These results demonstrated that PLR was significantly higher in patients with VTE. A PLR value of > 320 was found to have an independent and strong association with the occurrence of VTE (β 95% CI: 5.3256 - 5.3868; p < 0.0001). In addition to PLR, length of surgery was also found to have an independent association with VTE (β 95% CI: 0.0001 - 0.0006; p = 0.0056). The results of our analysis are consistent with published reports in other cancers in the ambulatory setting by Yang^13^, and Ferroni.^12^ These authors showed that pretreatment PLR was associated with VTE in the ambulatory chemotherapy setting. In this paper, we have shown that this association is also present in the inpatient postoperative setting, for HNC patients.

Although the results of this analysis are positive, the exact underlying mechanisms behind the observed association of PLR and thrombosis are speculative. ^10–13^ In the past few years, there has been increasing interest in the tumor microenvironment and its effect on systematic inflammation. ^28^ This has led to the study of other biomarkers in addition to PLR, ^29, 30^ such as the neutrophil-to-lymphocyte ratio (NLR)^29, 30^ and the lymphocyte-to-monocyte ratio (LMR) ^31^ in HNC prognosis. The ‘seed and soil’ hypothesis postulates that the tumor extracellular matrix (ECM) interacts with and is in turn affected by the inflammatory cells and mediators. ^32, 33^ Kuss and colleagues had previously demonstrated that there is altered lymphocyte homeostasis in SCC of the head and neck, resulting in reductions in all T-cell subsets. ^34^ Since malignancies have been known to be associated with thrombotic events, ^35, 36^ the lymphocyte count might thus be acting as a nonspecific surrogate for magnitude of the tumor’s systemic interaction – and by extension, its thrombotic potential.^12, 13^

On the other hand, the link between platelets and cancer-associated thrombosis has been well documented. ^37–39^ By combining the platelet and lymphocytes counts into the PLR, a crude ‘summary effect’ between the tumor-induced prothrombotic (platelet) and the inflammatory states (lymphocyte) is encapsulated. The superiority of the PLR over platelet count or WBC count alone was demonstrated in our study, in agreement with other authors. ^12, 13^ Ferroni and colleagues speculated that increased levels of platelets in itself could be related to underlying inflammation. In this context, our results were also in agreement, showing a small but highly significant relationship between the platelet count and lymphocyte count (R^2^ = 0.05, p < 0.0001), and platelet count and WBC (R^2^ = 0.08, p < 0.0001).

Consistent with other studies, our results also showed that length of surgery was also an independent predictor of VTE. ^40^ The association of length of surgery, though highly significant, was not shown to be as strong as PLR, demonstrated by the marginal β coefficient. Our results did not show postoperative heparin or aspirin to be associated with VTE. This could be due to the high amounts of patients already receiving postoperative VTE chemoprophylaxis in this high-risk cohort – which would therefore require a much larger sample size to detect a significant difference.

There are limitations to our study that we acknowledge. Our study was conducted as a retrospective cohort study, and as such, is subject to the intrinsic drawbacks of all retrospective data collection studies. Secondly, our study had a small sample size with small number of VTEs. This problem of small sample sizes is a similar problem faced by other authors investigating VTEs in HNC, with Thai et al and Clayburgh et al reporting 2 and 8 VTEs respectively. ^6, 9^ Unlike other published studies of VTE in the HNC literature, we compensated for our small data set with statistical techniques utilizing bootstrap ^22^ and statistical penalization. ^19^

The results reported here have shown that PLR has a significant and strong association with VTE. Current guidelines by ASCO ^15^ already recommend VTE chemoprophylaxis for patients with malignant disease undergoing surgery. In the study authored by Clayburgh et al, ^9^ they prospectively performed ultrasound scans (USS) for all postoperative patients and found that the incidence of VTE was much higher than reported in previous retrospective studies. Drawing from this result, they hypothesized that a significant proportion of asymptomatic or mildly symptomatic VTEs go undetected in patients with HNC. Furthermore, Thai and Clayburgh also showed that above the highest Caprini risk level, there was a failure to discriminate which patients would eventually develop a VTE. ^6, 9^ Thus, it is possible that the future utility of the PLR may be to stratify within high-risk HNC patients - to decrease the clinical threshold in which to perform diagnostic VTE tests in patients with elevated PLR.

## Conclusion

This exploratory pilot study has shown that PLR offers a potentially accurate risk stratification measure as an adjunct to current tools, without additional cost to health systems. Future large prospective studies are needed in order to fully delineate the characteristics of the relationship between the PLR and thrombotic risk.

## Acknowledgements

We would like to thank Yonatan Bardash, BS (Hofstra-Northwell School of Medicine, New York) for assistance in building our REDCAP database. We would also like to thank our statistical consultants, Lydia Hsu, PhD (Columbia University, Department of Statistics, New York), and Guillaume Stoffels, MS (Feinstein Institute of Medical Research, Department of Biostatistics, New York) for their assistance with the statistical analyses.

